# Improving classification and reconstruction of imagined images from EEG signals

**DOI:** 10.1101/2022.06.01.494379

**Authors:** Hirokatsu Shimizu, Ramesh Srinivasan

**Affiliations:** Department of Cognitive Sciences, University of California, Irvine, CA, USA; Department of Biomedical Engineering, University of California, Irvine, CA, USA

## Abstract

Decoding brain activity related to specific tasks, such as imagining something, is important for brain computer interface (BCI) control. While decoding of brain signals, such as functional magnetic resonance imaging (fMRI) signals and electroencephalography (EEG) signals, during observing visual images and while imagining images has been previously reported, further development of methods for improving training, performance, and interpretation of brain data was the goal of this study. We applied a Sinc-EEGNet to decode brain activity during perception and imagination of EEG stimuli, and added an attention module to extract the importance of each electrode or frequency band. We also reconstructed images from brain activity by using a generative adversarial network (GAN). By combining the EEG recorded during a visual task (perception) and imagination task, we have successfully boosted the accuracy of classifying EEG data in the imagination task and improved the quality of reconstruction by GAN. Our result indicates that the brain activity evoked during the visual task is present in the imagination task and can be used for better classification of the imagined image. By using the attention module, we can derive the spatial weights in each frequency band and contrast spatial or frequency importance between tasks from our model. Imagination tasks are classified by low frequency EEG signals over temporal cortex, while perception tasks are classified by high frequency EEG signals over occipital and frontal cortex. Combining data sets in training results in a balanced model improving classification of the imagination task without significantly changing performance in the visual task. Our approach not only improves performance and interpretability but also potentially reduces the burden on training since we can improve the accuracy of classifying a relatively hard task with high variability (imagination) by combining with the data of the relatively easy task, observing visual images.

## Introduction

Decoding the contents of the mind from brain activity is the core task in designing and implementing brain computer interfaces (BCI). A common task for in BCI research is to decode imagined movements [1] in order to control the movements of an external device [2], e.g., a computer mouse or a wheelchair, and more recently to facilitate rehabilitation [3]. Such BCI are potentially useful in applications focused on direct interaction with the physical world. A more challenging goal is to use BCI technology to decode the contents of the mind with the goal of communication [4]. A common type of decoding BCI involves selecting one object out of many by detecting the focus of attention, as in a P300 speller [5] or SSVEP based BCI [6]. More recently, there have been a number of studies focused on decoding imagined speech [7, 8] from electroencephalography (EEG) to potentially allow for communication BCI.

Another potentially useful approach that has garnered recent interest is decoding visual imagery using functional magnetic resonance imaging (fMRI) signals [9] and EEG signals [10]. One EEG study used handcrafted time-frequency features extracted from EEG signals and achieved high classification accuracy (∼90%) in four way classification [10]. A number of studies have focused on decoding imagined geometric shapes using deep learning methods [11–14]. While simple geometric shapes are potentially useful, broader applicability of this approach would require identification of a broad range of imagined images.

In the classification of perceived images from EEG activity, very strong classification results have been obtained in studies very large numbers of image classes (40) combining data generated from single exposures to 50 different sample images to each class. The classification performance in these tasks are 20 ∼ 50% in different studies [15, 16]. These robust results have even motivated attempts to reconstruct the images from EEG activity during the perception of the image using a generative adversarial network (GAN) [17]. They have also motivated many aspects of the present study.

In this study, we attempt to answer 4 scientific questions, (1) are the brain networks for decoding perceived and imagined images similar, (2) can we boost the performance of decoding imagined and perceived images by jointly training networks, (3) can we develop methods to identify brain activity giving rise to decoding when using neural network classification, (4) is the decoding of imagined images sufficient to enable the use of GAN technology to reconstruct imagined images. Our results indicate that the brain activity evoked during presentation of visual stimuli is present during imagination of the stimuli and can be used for better classification of the imagined image, and for improving performance in reconstructing the image.

## Materials and methods

### Ethics Statement

This study was approved by the UCI IRB for Human Subjects Research. Written informed consent was obtained from each subject prior to participation in experiments.

### Image dataset

The image dataset is a 40-class subset of the ImageNet dataset [18]. We used the same 40 classes as [17]. The dataset consists of 50 images per class, a total of 2000 images. Each image was cropped to the center with a size of 370 pixels by 370 pixels.

### Experiment

Four healthy subjects (1 female) participated in the experiment. EEG was recorded by a 128-channel NeuroScan EEG system with 2 kHz sampling rate. The experiment was conducted in a darkened room and each subject sat in a chair positioned in front of a desk and a monitor. Chin rest was not used in the experiment.

#### Visual experiment

In the visual experiment, the subjects were shown 40 blocks of a burst of 50 images of the same object class (500 ms per image, total of 25 seconds per block). The size of the image was 10 cm by 10 cm and the order of presentation was shuffled for each subject. Each block started with a fixation cross and ended with a blank black screen. The subjects were able to take a rest between each block and begin the next block when they were ready.

#### Imagination experiment

In the imagination experiment, each subject was shown one of the 40 images (1 image per class), which were picked from the dataset used in the visual experiment, for 500 ms and told to imagine the image with eyes opened. Each experiment block started with showing a gray square with a size of 20 cm by 20 cm as a canvas. Then a fixation cross was shown in the center of the gray square. One of the images was shown 500 ms followed by a 500 ms mask (noise image) with a size of 10 cm by 10 cm to flush buffered activity in the brain. After the noise image disappeared, the subjects tried to imagine the image on the gray square for 10 seconds. After 10 seconds of imagination, a blank black screen was shown. As in the visual experiment, the subjects were able to take a rest between each block and begin the next block when they were ready.

### Data preprocessing

The recorded EEG signals from both visual and imagination experiments were filtered by a band-pass filter with frequency of 1-50 Hz. For the visual experiment, the data was divided into 500 ms epochs corresponding to the presentation of each image. For the imagination experiments each 10 second recording was split into 20 trials of 500 ms without overlap.

As for the signals from visual experiments, trials with big distortion due to head or electrode movement were removed by visual inspection before removing ocular artifacts. Ocular artifacts in all signals were removed by deleting the ICA [19] components with high correlation to the EOG signal recorded by channels Fp1 and Fp2.

After removing artifacts, all trials were downsampled to 250 Hz. After downsampling, each trial was converted to the average reference and each channel was independently z-scored. In total, 7987 trials from visual experiments and 3200 trials from imagination experiments were obtained.

### Model

#### EEG classification model

In an EEG classification task, we used a model with Sinc-EEGNet architecture which contains Sinc-based convolution layer, depth-wise convolution layer, and separable convolution layer [20]. We used 16 filters with a kernel size of 65, 1 Hz for minimum frequency and 4 Hz for minimum bandwidth of the filters in the Sinc-based convolution layer, and 2 for a multiplier parameter in the depth-wise convolution layer. Also, we added 2 dense layers (512 and 2048 output size) between the flatten layer and the final dense layer with output size of 40 to make the model visual-guided [21]. A detail of the structure of the model is shown in Table 1.

**Table 1.** Architecture of the model.

| Layer | Parameters |
| --- | --- |
| Input | Shape: (128, 125, 1) |
| Sinc Convolution | Num of Filters: 16, Kernel Size: (1, 65),<br>Freq Min: 1.0, Band Min: 4.0 |
| Batch Normalisation |  |
| Depth-wise Convolution | Kernel Size: (128, 1), Multiplier: 2 |
| Batch Normalisation |  |
| Activation | Elu |
| Average Pooling | Size: (1, 4) |
| Dropout | Rate: 0.5 |
| Separable Convolution | Num of Filters: 32, Kernel Size: (1, 8) |
| Batch Normalisation |  |
| Activation | Elu |
| Average Pooling | Size: (1, 3) |
| Dropout | Rate: 0.5 |
| Flatten |  |
| Dense | Size: 512, Activation: Relu |
| Dropout | Rate: 0.5 |
| Dense | Size: 2048, Activation: Relu |
| Dropout | Rate: 0.5 |
| Dense | Size: 40, Activation: Softmax |

The loss function for this model is a summation of softmax cross-entropy loss and mean squared error of the extracted feature, as shown in the Eq (1).

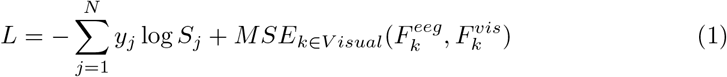

In the equation, *S*_*j*_ is the softmax output of category *j, y*_*j*_ is the label of input, and *N* is the number of classes. The second term is a mean squared error between the features 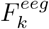 extracted from EEG signals and the features 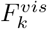 extracted from the corresponding image, which is the output of the penultimate layer (whose output size is 2048) of the ResNet101 [22]. The second term was calculated only if the EEG signals were from visual experiment.

#### Attention mechanism

To study which frequency band and electrodes are important for classification, we added an attention module to the model, by adding a squeeze-and-excitation module [23] after the batch normalization layer following the depth-wise convolution layer. When applying the squeeze-and-excitation module, we used both average-pooled features and max-pooled features [24]. The diagram of the attention module is shown in Fig 1. The shared network has one hidden layer with size of 32*/r*, where *r* is the reduction ratio.

**Fig 1.**
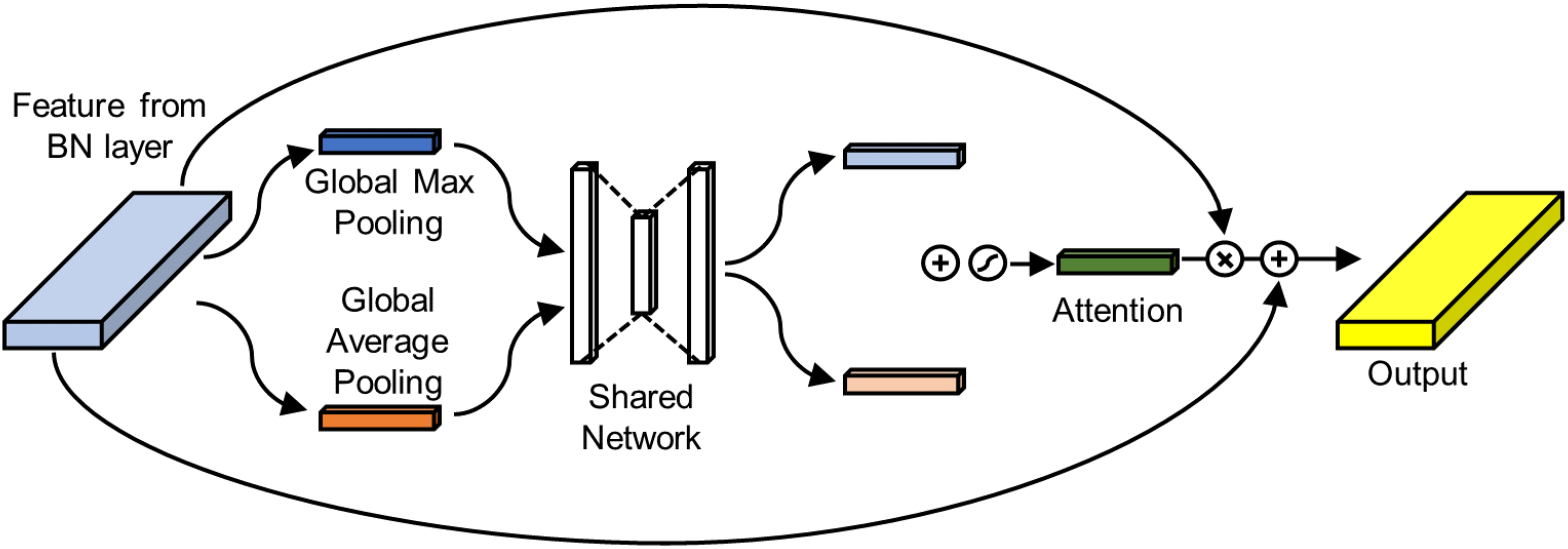
Diagram of the attention module. As illustrated, an element-wise summation of each output from the shared network, a sigmoid function activation, an element-wise multiplication between attention and initial features, and an element-wise summation with initial features are performed after the shared network.

#### GAN

We used a GAN framework which uses features extracted from EEG signals to condition a generative network to visualize the seen image in the visual experiment and the imagined image in the imagination experiment. The EEG features are the outputs of the dense layer whose output size is 2048 in the EEG classification model. The architecture of the GAN is inspired by recent work [25].

The generator takes EEG features with 2048 size and random noises with 2048 size from a normal distribution as input. After the input layer, we added a trainable weighted Gaussian layer [26]. The trainable weighted Gaussian layer has two trainable parameters, mean (*μ*) and variance (*σ*), and translates the input noises (*ϵ*) to a mixture of Gaussian distributions and then uses EEG features (*eeg*) as weights. The output of this layer is shown in Eq (2).

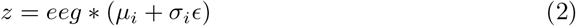

After the weighted Gaussian layer, the input is fed into 5 transposed convolution blocks. The first 4 blocks contain a transposed convolution layer, batch normalization layer and LeakyRelu activation layer, and the final block contains a transposed convolution layer and tanh activation layer. The detailed architecture of the transposed convolution layer is shown in Table 2.

**Table 2.** Architecture of the generator.

| Block | Filters | Kernel Size | Strides | Padding |
| --- | --- | --- | --- | --- |
| 1 | 512 | 4 | 1 | valid |
| 2 | 256 | 4 | 2 | same |
| 3 | 128 | 4 | 2 | same |
| 4 | 64 | 4 | 2 | same |
| 5 | 3 | 4 | 2 | same |

The architecture of the discriminator is inspired by VGG16 [27] and contains 10 convolution blocks. Each block contains a convolution layer, batch normalization layer and LeakyRelu activation layer. The max pooling layer, which halves the map size, is applied after the second, the fourth, the seventh and the last blocks. After the final convolution block, the output is flattened and fed into two classifiers which distinguish real images and classify the image category. The binary classifier for distinguishing real images has two dense layers whose output sizes are 1024 and 1. The activation function is Relu for the first dense layer and sigmoid for the second layer. The multi-class classifier for classifying the image category has three dense layers whose output sizes are 1024, 1024 and 40. The activation function is Relu for the first and second dense layer and softmax for the third layer. Additionally, the dropout layers are added after each max pooling layer and dense layer.

The loss function for the discriminator is a summation of binary cross-entropy loss for identifying real images as real and generated images as fake, and softmax cross-entropy loss for classifying the real images. The loss function for the generator is a summation of binary cross-entropy loss for identifying generated images as real, and softmax cross-entropy loss for classifying the generated images.

## Result

We split the dataset into training (80%), validation (10%) and test (10%) sets. For the visual experiment, we split the paired image-EEG data (0.5 s) into training, validation, and test sets [28]. As for the imagination experiment, each period of imagination (10 secs) was divided into labeled segments matching the segment lengths (0.5 s) of the visual experiment. This labeled imagination dataset was divided into training, validation, and test sets for each image label.

### Classification accuracy

Model training was performed by using Adam optimizer with learning rate of 5e-4. We used an early stopping strategy with patience of 30 epochs to prevent overfitting. We trained the model in three ways, using the dataset from visual experiment (“Visual”), using the dataset from imagination experiment (“Imagine”), and using the dataset that combines the dataset from visual and imagination experiments (“Mix”). The batch size for each training was 128 for “Visual” and “Mix”, 64 for “Imagine”, which was adjusted to be about the same number of steps for each epoch.

Fig 2 shows the performance on the validation set. The curve of the “Mix” model shows the accuracy of classifying the visual validation set in Fig 2A, and the imagination validation set in Fig 2B. Chance accuracy is 0.025 (1/40) and is shown by the dashed gray line. Table 3 shows a summary of the accuracy of classifying the imagination test set. To calculate an averaged accuracy, we trained the model by using the same training, validation and test sets 10 times. To compute performance for individual subjects, we only used each subject’s dataset for training, validation, and testing of the “Imagine” model, and combined each subject’s dataset with all subject’s dataset from visual experiment for the “Mix” model.

**Fig 2.**
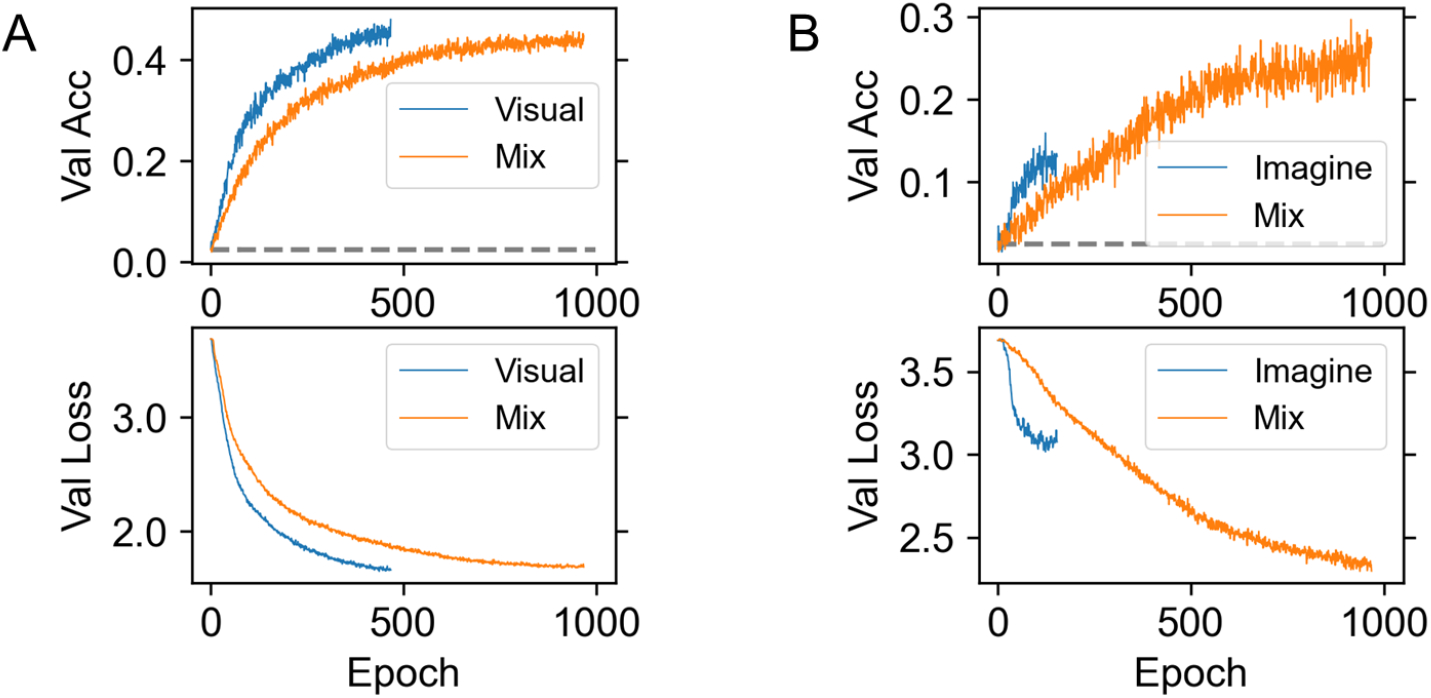
Validation curve. (A) Performance on the validation set of the visual experiments. (B) Performance on the validation set of the imagination experiments. The chance accuracy is 1/40 = 0.025 (shown by the gray line).

**Table 3.** Averaged accuracy of classifying the imagination test set.

|  | Imagine | Mix |
| --- | --- | --- |
| Subject 1 | 0.211 ± 0.021 | 0.334 ± 0.034 |
| Subject 2 | 0.176 ± 0.020 | 0.260 ± 0.045 |
| Subject 3 | 0.094 ± 0.035 | 0.155 ± 0.041 |
| Subject 4 | 0.132 ± 0.058 | 0.249 ± 0.037 |
| All | 0.134 ± 0.042 | 0.252 ± 0.022 |

Test accuracy was improved for both each subject’s result and combined result when we used the “Mix” model.

The accuracy of classifying the imagination data can be improved by using a training dataset that combines the visual data and the imagination data. The accuracy of classifying the visual data is comparable when comparing training with just the visual data and the combined visual and imagination data. This result suggests that the brain activity evoked in the visual experiment is present in the imagined experiment and can be used to better classify the imagined image. The neural network does not select this activity for classification when using the imagined data alone for training and thus does not perform as well. We thus focused on using an attention methods to understand the nature of the difference in brain activity used by these models.

### Attention

A model with an attention module was trained in the same way as the model without an attention module. We used 8 as a reduction ratio (we also performed the training with a reduction ratio of 4, and we obtained results with similar trends.).

Fig 3 shows frequency bands of spatial filters sorted based on an attention map averaged over the test set. For example, “Top 1” means the filter with the highest attention value. For the “Mix” model, the averages of attention maps were calculated separately for visual test set (“Mix (Visual)”) and imagination test set (“Mix (Imagine)”). As can be seen from Fig 3, the Sinc-based convolution layer creates more wide band filters at high frequency regions and pays higher attention to them when trained by the visual dataset (“Visual”) than trained by the imagination dataset (“Imagine”). Also, we can see that the “Mix” model pays attention to both high and low frequency bands compared to the “Visual” or “Imagine” model. This means that the “Mix” model produces a balanced result of visual and imagination dataset even though the size of the dataset is not balanced, and wide bands at high frequency regions, which are not created when using only imagination dataset, improve the accuracy of the imagination classification.

**Fig 3.**
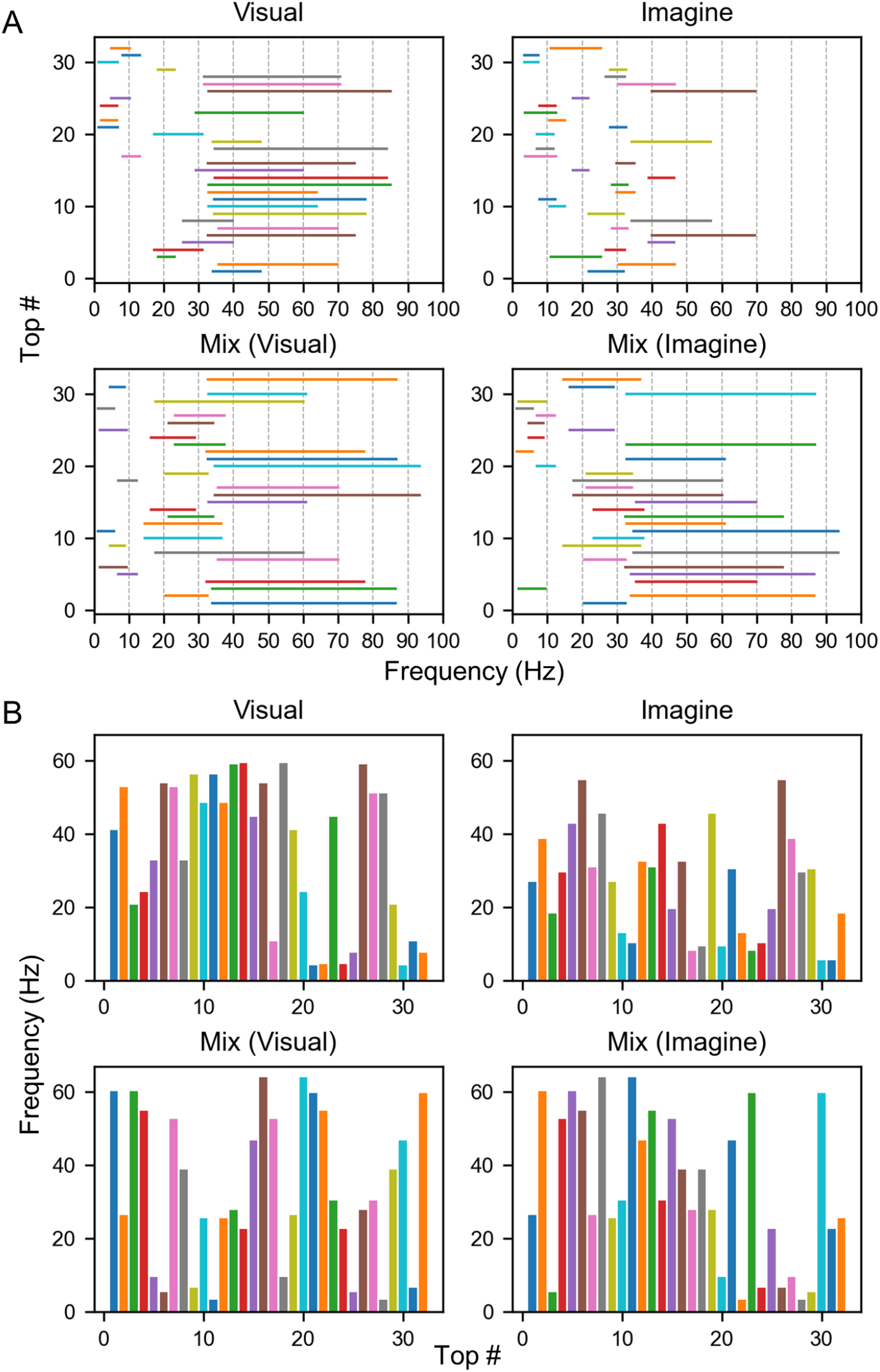
Sorted frequency band of each model. Sorting was performed based on the attention map averaged over the test set. “Top 1” means the filter with the highest attention value. (A) Frequency band of each spatial filter. (B) Center frequency of each spatial filter.

To analyze the spatial filters, we calculated the weighted sum of absolute values of spatial filters as shown in Fig 4A. We took an absolute value because we considered that position with large value in the filter is more important regardless of the sign of those values. Also, we used the averaged attention map as weights because spatial filters with higher attention value mean that those filters are more important. The map was normalized so that the maximum value is 1. We calculated the weighted sum of sinc functions and performed the Fourier transform to represent the spectral composition of the filters as shown in Fig 4B.

**Fig 4.**
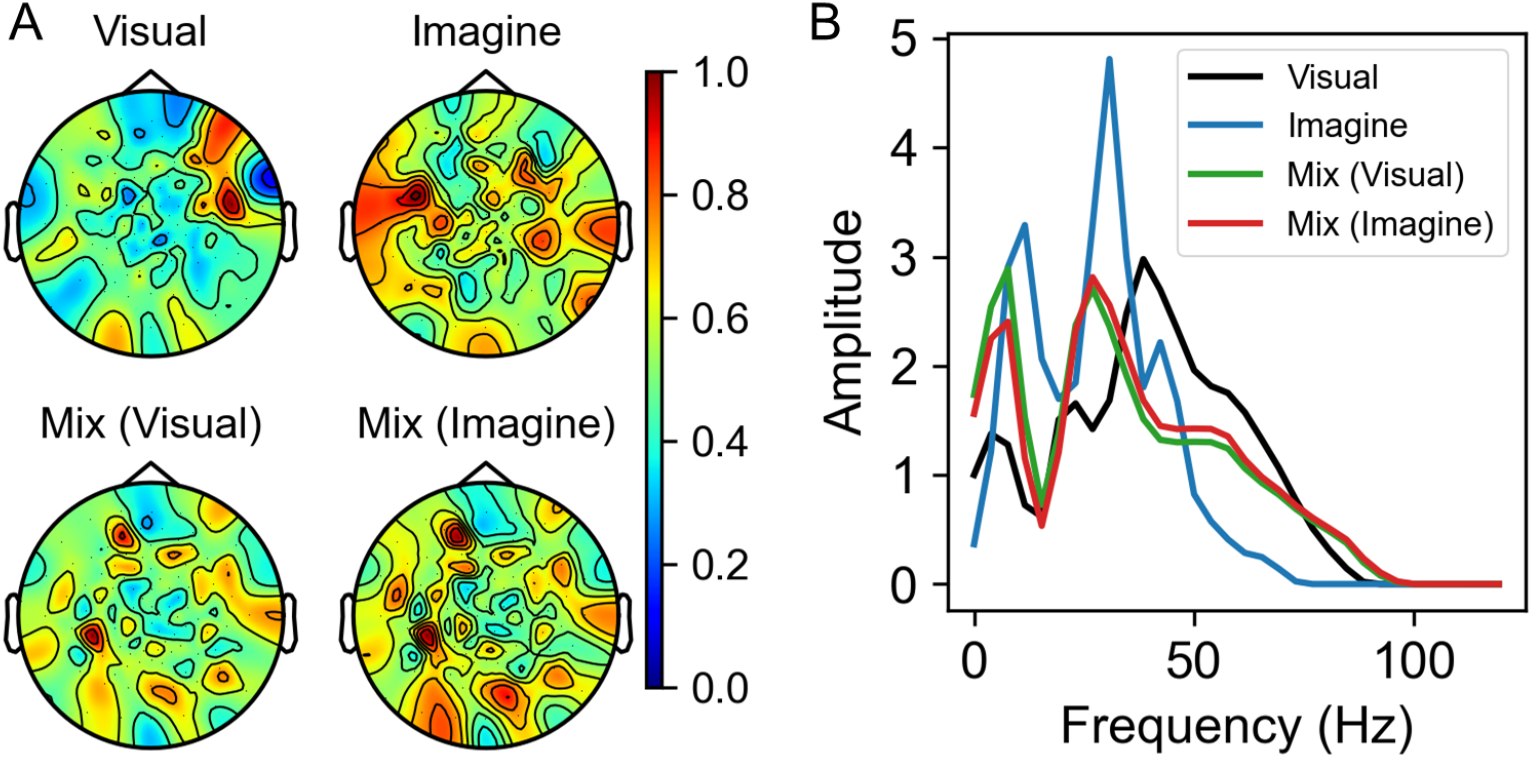
Weighted sum results. (A) Weighted sum of absolute values of spatial filters. Each result was normalized so that the maximum value is 1. (B) FFT results of weighted summed sinc filters.

As can be seen from Fig 4A, the spatial filters in the “Visual” model are weighted to the occipital and frontal lobes, while the “Imagine” model is weighted towards the temporal lobe. In the “Mix (Visual)” model get relatively higher weights in the temporal lobe area than the filters in the “Visual” model and in the “Mix (Imagine)” model more weight is placed on the occipital and parietal lobe area than the filters in the “Imagine” model. This can also be confirmed from Fig 5, which shows the difference between each weighted summed spatial filter. Fig 4B shows that the “Mix (Visual)” model has higher weight in the low frequency region than the “Visual” model and the “Mix (Imagine)” has higher amplitude in the high frequency region than the “Imagine” model. These results also indicate the “Mix” model produces a balanced weighting of visual and imagination brain activity thereby improving performance.

**Fig 5.**
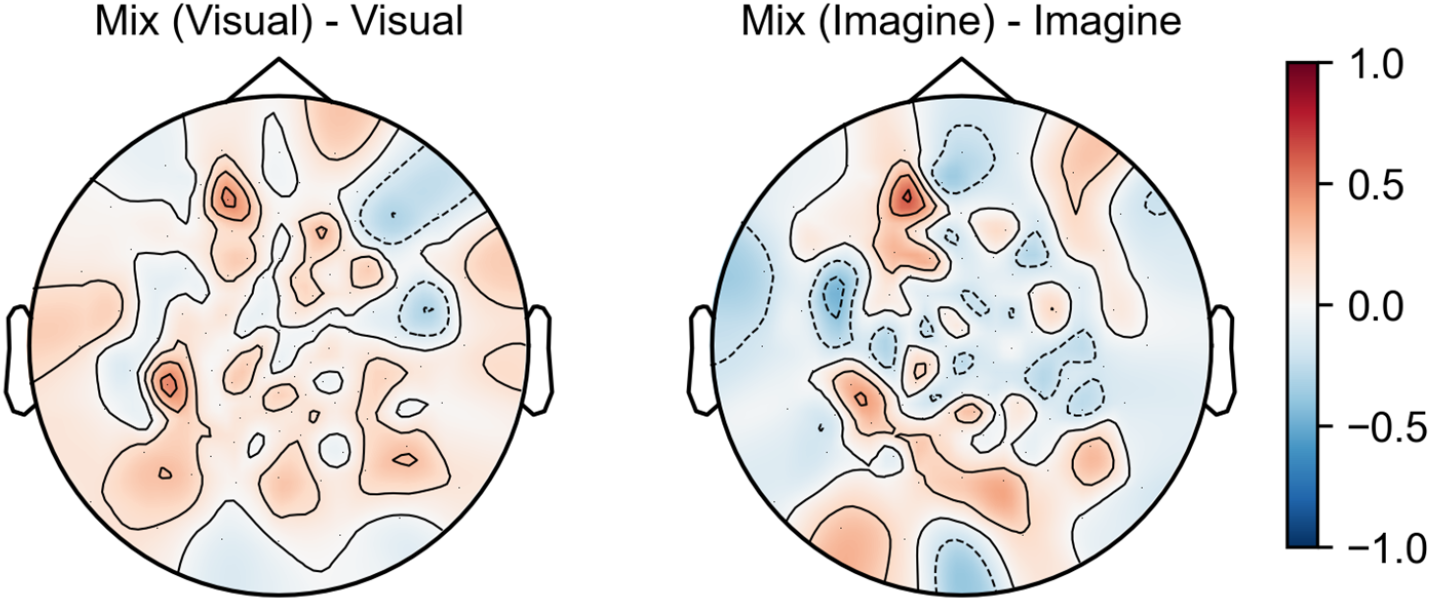
Difference between filters. The map of the difference between the weighted sums of the spatial filters in the “Mix” model and the “Visual” model which were derived from the visual test set (Left), and between the “Mix” model and the “Imagine” model which were derived from the imagination test set (Right).

We divided the frequency ranges in Fig 3, we can see 3 blocks of frequency, “Low” (alpha/mu band or below, *<* 15 Hz), “Mid” (beta band, 15 - 35 Hz), and “High” (gamma band, *>* 35 Hz). For a more detailed analysis, we manually picked filters in each frequency block and calculated the weighted sum of filters. Table 4 shows numbers of picked filters in each block. The numbers correspond to “Top #” in Fig 3. Fig 6 shows the results of weighted summed filters in each block. Each map was normalized so that the maximum value is 1. We can see that the “Low” block has large values in the occipital area and the “High” block has large values in the frontal area.

**Table 4.** Numbers of picked filters in each frequency block.

|  | Low | Mid | High |
| --- | --- | --- | --- |
| Visual | 17, 21, 22, 24, 25, 30, 31, 32 | 3, 4, 5, 8, 20, 29 | 1, 2, 5, 6, 7, 8, 9, 10, 11, 12, 13, 14, 15, 16, 18, 19, 23, 26, 27, 28 |
| Imagine | 10, 11, 17, 18, 20, 22, 23, 24, 30, 31 | 1, 3, 4, 7, 9, 12, 13, 15, 16, 21, 25, 28, 29, 32 | 2, 5, 6, 8, 14, 19, 26, 27 |
| Mix (Visual) | 5, 6, 9, 11, 18, 25, 28, 31 | 2, 8, 10, 12, 13, 14, 19, 23, 24, 26, 27, 29 | 1, 3, 4, 7, 8, 15, 16, 17, 20, 21, 22, 29, 30, 32 |
| Mix (Imagine) | 3, 20, 22, 24, 26, 27, 28, 29 | 1, 7, 9, 10, 14, 16, 17, 18, 19, 25, 31, 32 | 2, 4, 5, 6, 8, 11, 12, 13, 15, 16, 18, 21, 23, 30 |

The numbers correspond to “Top #” in Fig 3. If the frequency band of a specific filter spans multiple blocks, the filter is used for each block.

**Fig 6.**
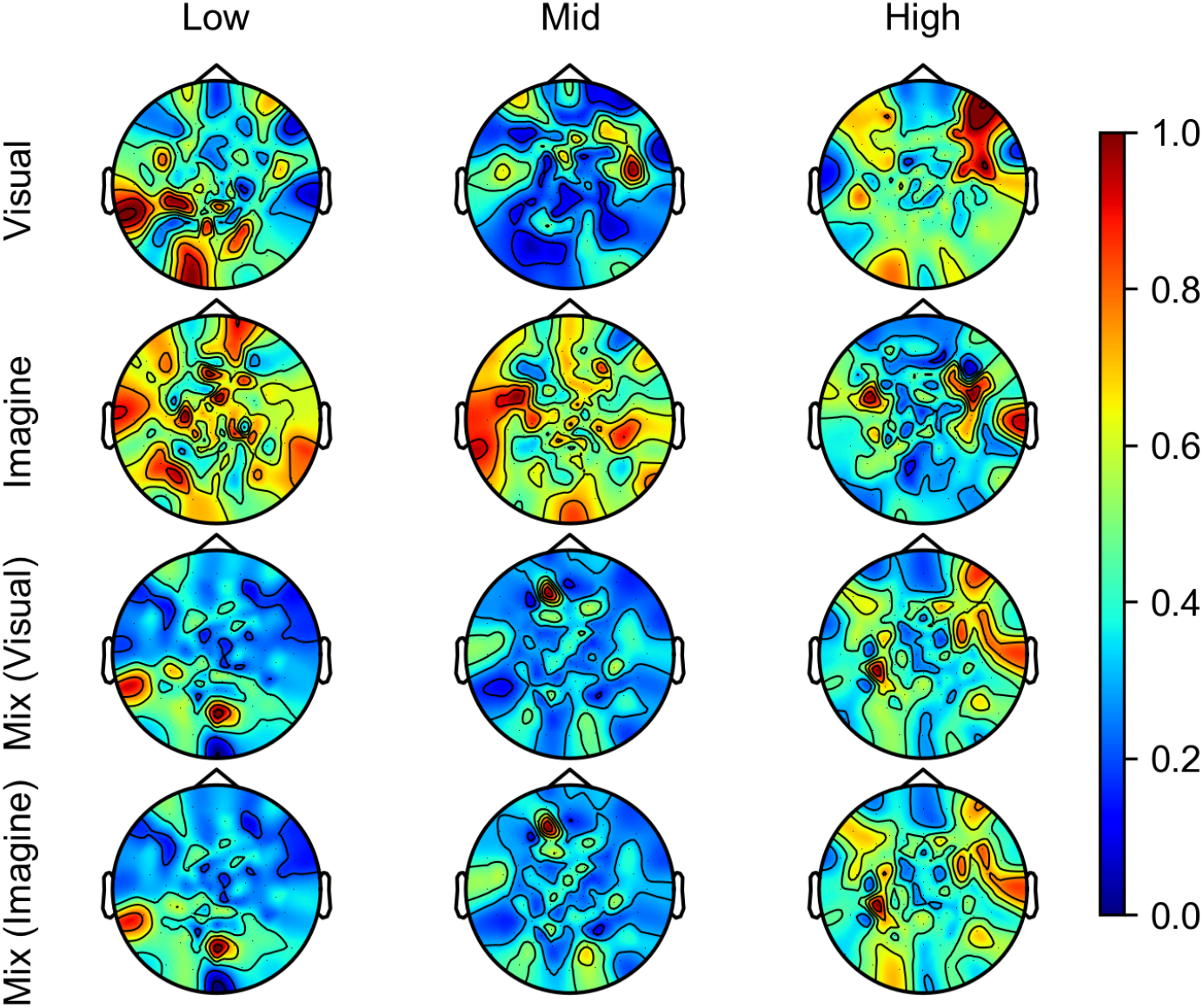
Weighted summed filters in each frequency block. Each map was normalized so that the maximum value is 1. “Low” block mainly contains alpha band or below, “Mid” block mainly contains beta band, and “High” block mainly contains gamma band.

### GAN

We trained a GAN for 1000 epochs by using training and validation sets of the visual experiments as a training dataset. Adam optimizer with learning rate of 1e-4 and with *β*_1_ of 0.5 was used for training. The batch size was 16.

To evaluate the generated images, we made an image classifier trained on the images from the same 40-class subset of the ImageNet dataset which contains all images other than those used in EEG classification, a total of 49425 images. The architecture of the image classifier is the same as ResNet101 except that the final output size is changed to 40 (originally 1000). We used the ResNet101 model pre-trained on ImageNet dataset, and we additionally trained only the final output layer’s weights for 50 epochs.

“Visual” GAN is the GAN which is trained by using conditions extracted from the “Visual” classifier, and “Mix” GAN is the GAN which is trained by using conditions extracted from the “Mix” classifier. Fig 7 and 8 show samples of images generated from training (training and validation sets) or test dataset of the visual experiment, and the map of the image classifier’s accuracy. The image categories are dog, cat, butterfly, sorrel, capuchin, elephant, panda, fish, airliner, broom, canoe, cellphone, mug, convertible, computer, watch, guitar, locomotive, espresso maker, chair, golf ball, piano, iron, jack-o’-lantern, mailbag, missile, mitten, bike, tent, pajama, parachute, pool table, radio telescope, camera, gun, shoe, banana, pizza, daisy and bolete from the upper left. The generated images were then classified with ResNet with overall accuracy of 0.331 for the training set and 0.183 for the test set (chance is 0.025) for the “Mix” GAN. Some images (e.g., panda, elephant, fish) were identified with very high accuracy in both train and test sets, but others were only identified in the training set by ResNet, despite being equally obvious to a human observer (e.g., daisy, pumpkin), suggesting some of the performance limitations reflect the capabilities of ResNet.

**Fig 7.**
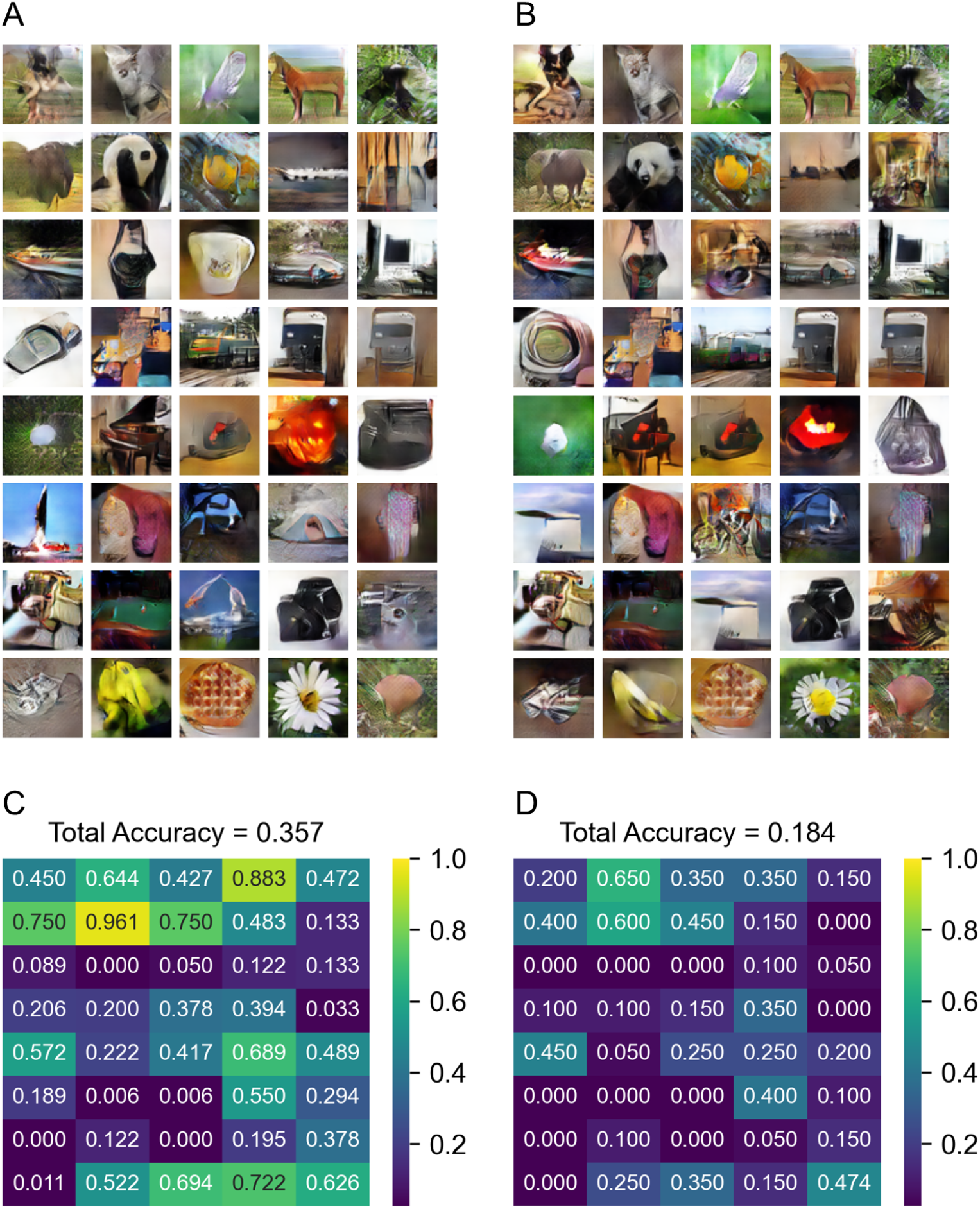
Results generated by “Visual” GAN. (A) Sample of images generated from the visual train dataset. (B) Sample of images generated from the visual test set. (C) The classification accuracy map of images generated from the visual train dataset. (D) The classification accuracy map of images generated from the visual test set.

**Fig 8.**
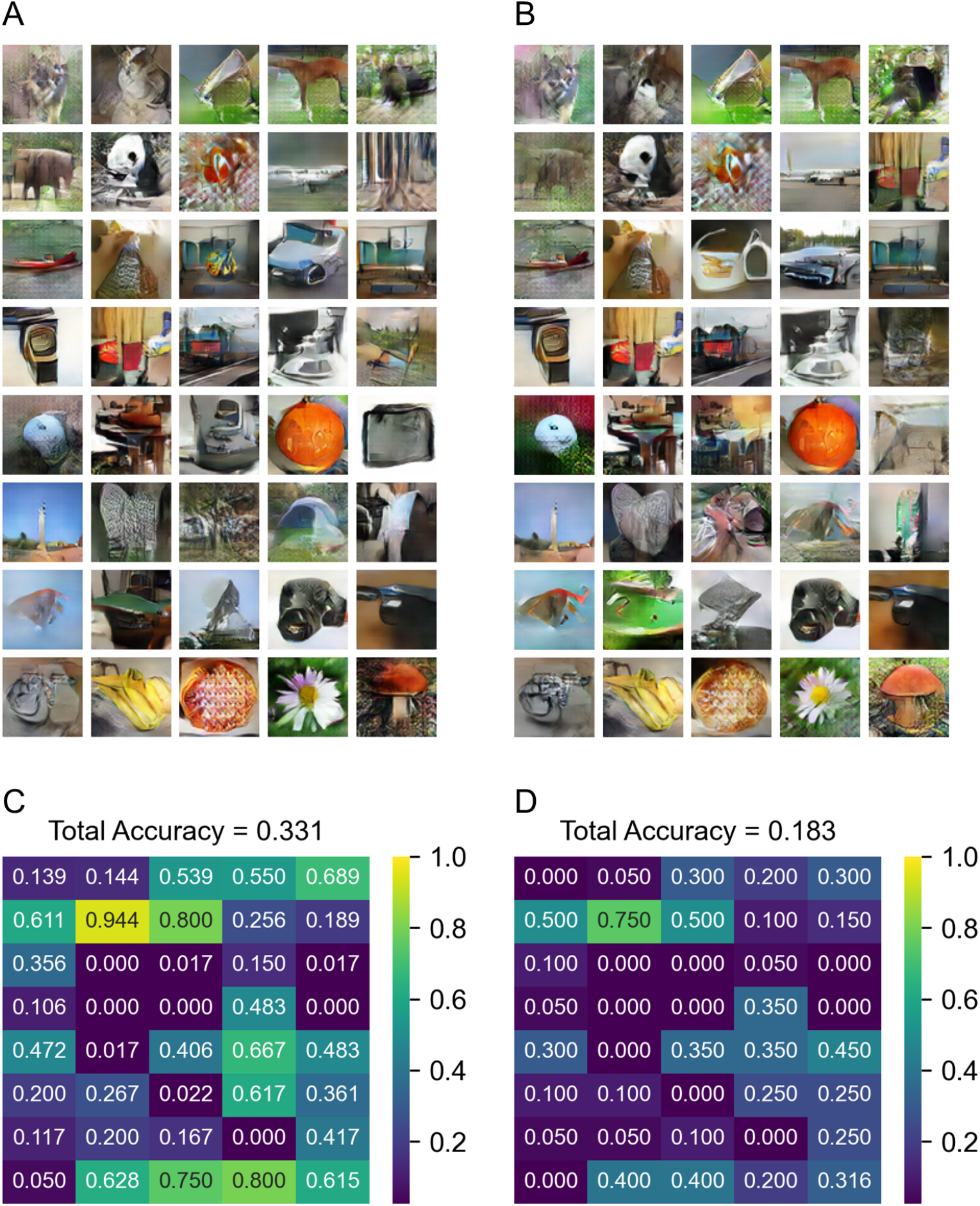
Results generated by “Mix” GAN. (A) Sample of images generated from the visual train dataset. (B) Sample of images generated from the visual test set. (C) The classification accuracy map of images generated from the visual train dataset. (D) The classification accuracy map of images generated from the visual test set.

Fig 9 shows the results generated from the imagination dataset (include all data). The performance of the “Visual” GAN is very poor on the imagination data. Since the imagination dataset is used when training the “Mix” classifier and the model is trained to use information from imagination data, some images can be generated better when using the “Mix” GAN, even though the imagination dataset is not used for training the GAN. The performance of the ResNet classifier is slightly above chance, and certain images (butterfly, daisy, pizza, pumpkin, etc.) can be generated by the GAN and identified by ResNet.

**Fig 9.**
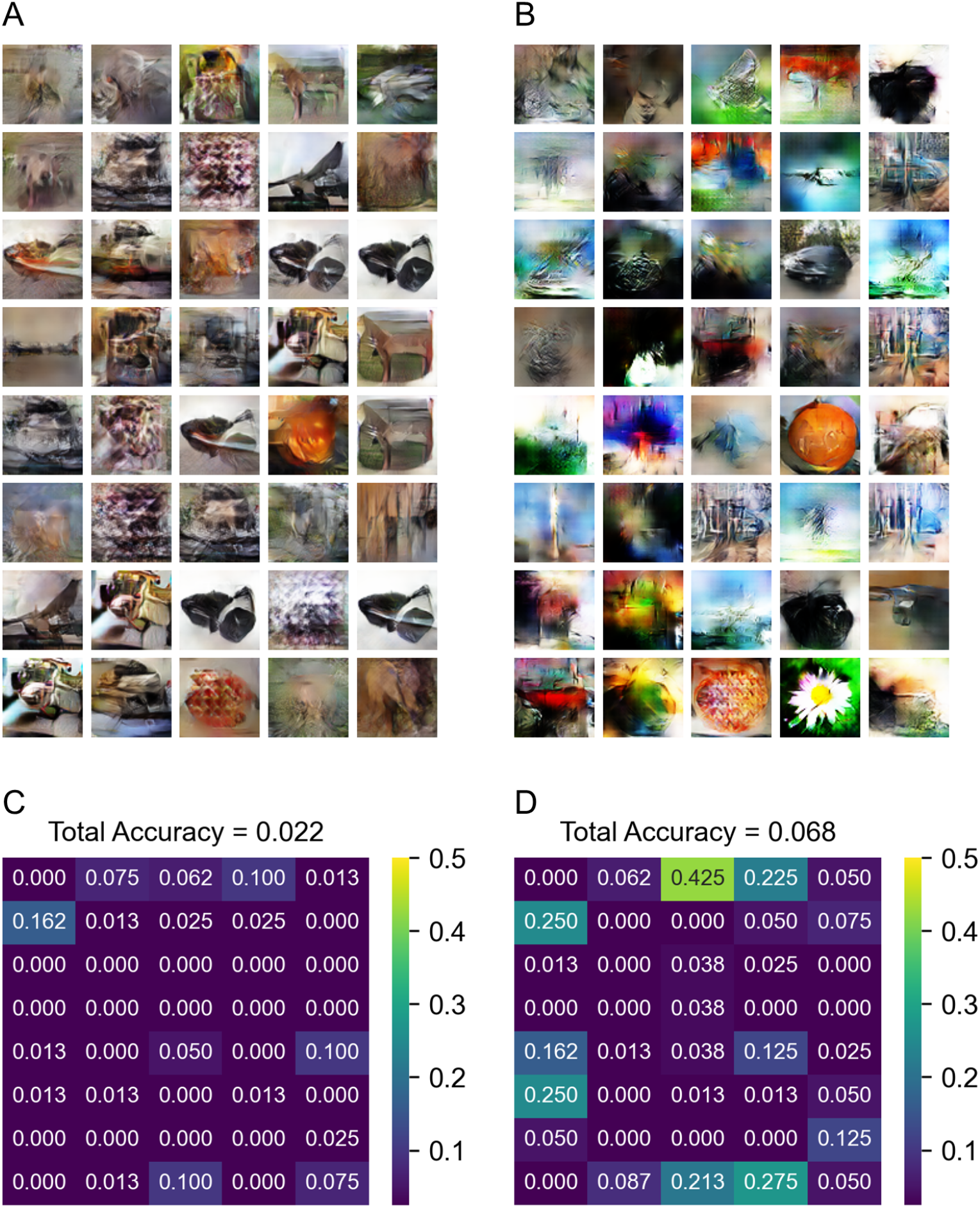
Results generated from imagination dataset. (A) Sample of images generated by the “Visual” GAN. (B) Sample of images generated by the “Mix” GAN. (C) The classification accuracy map of images generated by the “Visual” GAN. (D) The classification accuracy map of images generated by the “Mix” GAN.

## Discussion

We have presented a novel approach to improve the accuracy of classifying imagined images. By combining the data of a visual presentation experiment with data from an imagination experiment, we have successfully boosted the classifying accuracy of imagination data, and this trend was similar for each subject. Our method can not only improve the accuracy but also reduce the burden on the subjects because we can improve the accuracy of classifying the relatively hard task (imagination) by combining the data of the relatively easy task, observing visual images.

By using Sinc-EEGNet and the attention module, we have derived the spatial (electrode) and frequency importance which offered us more interpretability. We showed that the “Mix” model can create a balanced model and that wide bands at high frequency regions (lower and upper gamma band) improve the imagination classification which will not be achieved if we used only imagination data to train the model. We also showed that trained models pay more attention to the occipital area in low frequency range and the frontal lobes in high frequency range. By incorporating the attention module, we can investigate not only the importance of each electrode and frequency range but also their relationship and the order of importance.

We also have shown a new GAN-approach for reconstructing the imagination with images. By using conditions extracted from the visual dataset using the “Mix” model, we can train a GAN which can generate images from the imagination dataset. Our approach uses only data from visual experiments which have a stronger relationship between EEG signals and images, that is suitable for training the GANs, than the data from imagination experiments which are relatively difficult to associate EEG data with the specific images. It is generally difficult to train a GAN enough to generate an image that allows the classifier to be classifiable as can be seen in Fig 7 and 8, which show that images in some categories were not generated at all even from the visual training dataset. With that in mind, we can see a decent improvement of image generation from the “Visual” model (Fig 9A and C) to the “Mix” model (Fig 9B and D) in some of the categories. The performance of the model was only evaluated by automatic recognition by a neural network. Our subjective impression is that a human observer would recognize even more of these images.

Our results demonstrate that the approach presented here is useful both for improving the performance of Brain-Computer Interfaces that classify and reconstruct imagined images and for enhancing the interpretability of neural network BCI models based on EEG data.

## Acknowledgments

The authors thank Qinhua Sun and Zhibin Zhou for help with data collection, and Khuong Vo for discussion about machine learning methods.

